# Novel instrumented insole algorithm accurately approximates plantar flexor loading

**DOI:** 10.1101/2019.12.20.885228

**Authors:** Todd J. Hullfish, Josh R. Baxter

## Abstract

Plantar flexor loading is critical for ambulatory function but there are few wearable solutions to monitor loading. The purpose of this study was to develop and validate a method to quantify plantar flexor loading using a commercially-available instrumented insole. Seven healthy young adults completed a battery of functional activities to characterize a range of plantar flexor loading which included single leg heel raise, step down, and drop jump as well as walking and running at comfortable speeds. Lower extremity trajectories were captured using motion capture and ground reaction forces were recorded with embedded force plates as well as the instrumented insole. Measurements of plantar flexor loading quantified by the instrumented insole were compared to ‘gold standard’ inverse dynamics. We found that the insole loading calculation of plantar flexor moment was accurate to within 4.2% on average compared to inverse dynamics across all activities. Additionally, the calculated wave forms were strongly correlated with inverse dynamics (R_xy_ > 0.928). Our findings demonstrate the utility and fidelity of a new method for measuring plantar flexor loading using a commercially available instrumented insole. By leveraging this new methodology, it is now feasible to prospectively track and eventually prescribe plantar flexor loading outside of the clinic to improve patient outcomes.

## Introduction

Plantar flexor loading is critical for ambulatory function throughout the lifespan [1,2] and in injured populations [3–5]. While inverse dynamics is the ‘gold standard’ for assessing joint moments, there are few wearable solutions to continuously monitor plantar flexor loading [6]. Therefore, the purpose of this study was to develop and validate a method to quantify plantar flexor loading outside of a traditional biomechanics lab. Based on our criteria that this technique must work on patients outside of a biomechanics laboratory with minimal training, we decided to use a commercially-available instrumented insole that offered wireless data logging. To accomplish this, we compared sagittal plane moments about the ankle derived from our insole implementation with moments calculated using inverse dynamics during a series of functional activities. We hypothesized that our method of measuring loads would be accurate within 10% of inverse dynamics.

## Methods

Seven healthy young adults (5 males, 2 females; 31 ± 4 years; BMI 24.1 ± 3.5) performed a battery of functional tasks after providing written informed consent in this IRB approved study. During these functional tasks, we acquired motion capture and force plate data to quantify plantar flexor loading and measured the interaction between the right foot and shoe using an instrumented insole. Briefly, subjects wore athletic shorts and lab standard running shoes (Air Pegasus, Nike, Beaverton, OR). Using a 12-camera motion capture system at 100 Hz (Raptor Series, Motion Analysis Corp, Santa Rosa, CA), we tracked the 3-dimensional trajectories of 28 retro-reflective markers that we placed on the lower extremities of each subject [7]. Briefly, we placed markers on bony-identifiable landmarks to scale constrained musculoskeletal models and additional markers on the lower extremity to track segmental kinematics during the functional tasks. We collected ground reaction forces at 1000 Hz using 3 embedded force plates (BP600900, Advanced Mechanical Technology, Inc., Watertown, MA). During the same test session, we also measured plantar loading of the right foot using a commercially available instrumented insole (Loadsol, Novel Electronics, St. Paul, MN, USA). The instrumented insole measured normal forces applied to 3 force sensing zones located under the rear-, mid-, and forefoot at 100 Hz (**Figure 1A**). We logged these data to a smart device (iPod Touch, Apple, Inc., Cupertino, CA) and synchronized the motion capture and instrumented insole data by matching peak foot loads collected from subjects as they swayed side to side.

**Figure 1.**
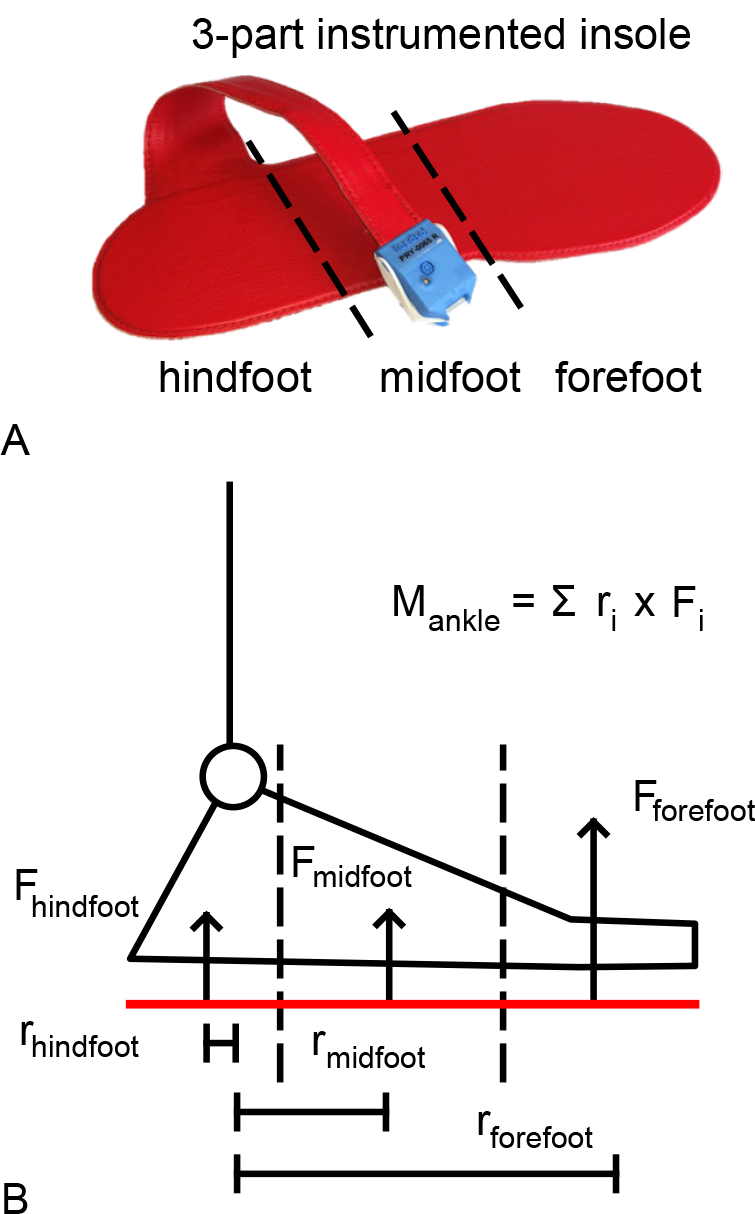
(single column): The instrumented insole is divided into 3 discrete zones, the hind-, mid-, and forefoot (A). The load applied to each of these zones occurs at a constant distance from the ankle joint center (B). As a result, the moment about the ankle can be quantified as the sum of the products of the moment arm and force applied to each zone.

Subjects performed a series of activities commonly used to evaluate plantar flexor function. We instructed participants perform single leg heel raises, step downs, counter movement jumps, and drop jumps as well as to walk and run at comfortable speeds. We selected these activities to characterize a range of plantar flexor loading magnitudes and rates as well as foot pitch angles. To quantify plantar flexor loading using a ‘gold standard’ technique, we used the motion capture and force plate data to solve the inverse dynamics problem at the right ankle joint. We scaled a generic musculoskeletal model to create subject specific models based on body weight and stature [7,8]. We then performed inverse kinematics and inverse dynamics to quantify plantar flexor loading during each activity.

To quantify the plantar flexor loading using the instrumented insole, we modeled each of the three force sensing zones as discrete one-dimensional force plates (**Figure 1B**). We assumed that the loads applied to each sensing zone were orthogonally directed. We also assumed that the centers of pressure of these forces were 50% of the anterior-posterior distance of the hindfoot and midfoot zones and 40% of the anterior-posterior distance of the forefoot zone. Because these centers of pressure locations do not change with respect to the ankle joint center, we applied a constant offset – measured as the distance from the most posterior aspect of the instrumented insole to the approximated ankle joint – to each of them to determine the moment arm of the load applied to each zone. Therefore, plantar flexor loading would be equal to the sum of the products of each zone moment arm and the applied load. To determine the fidelity of approximating plantar flexor loading using an instrumented insole, we calculated the peak loading errors, root mean square errors, and cross correlations, between the ‘gold standard’ inverse dynamics and our novel implementation of an instrumented insole.

## Results

We found that calculating peak plantar flexor loading using an instrumented insole was accurate to within 4.2% on average compared to the ‘gold standard’ inverse dynamics calculations across all activities (**Figure 2**). Additionally, the instrumented insole loading calculation produced waveforms that strongly correlated with motion capture approximations (R_xy_ > 0.928, **Figure 3**). Similarly, the root mean square errors for the movements averaged less than 0.6% body weight body heights, which is approximately 8.4 N m for a 1.8 meter tall, 82 kg person. High impact movements generated the largest errors during weight acceptance (10% of movement) and were still within 7% of motion capture derived plantar flexor moments.

**Figure 2.**
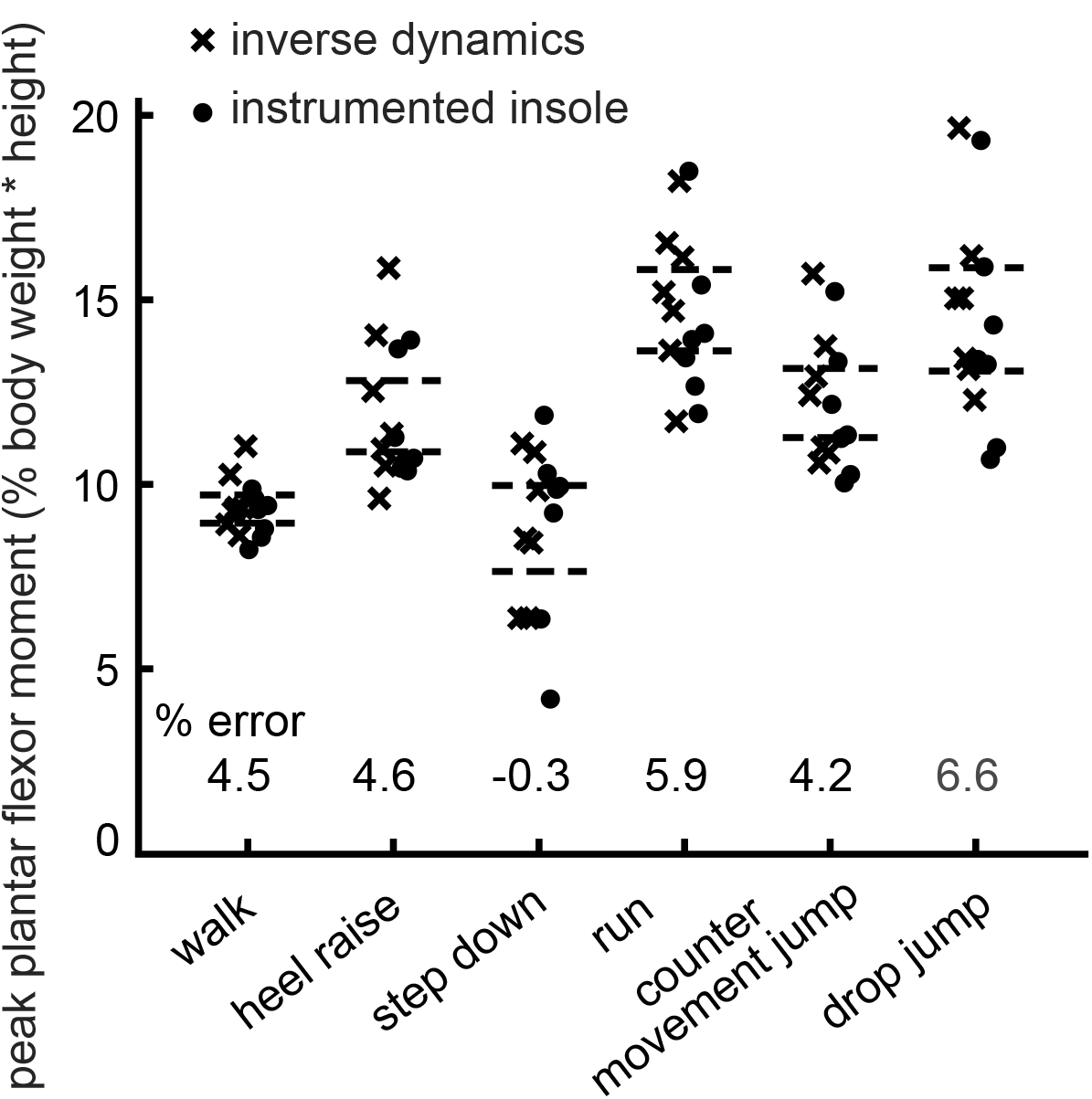
(single column): Measurements of peak plantar flexor loading from inverse dynamics (crosses) and the instrumented insole (dots) show good agreement across all activities we tested. The 95% confidence intervals for each measurement are represented with dashed lines. These measurements were accurate to within 4.2% on average with less than 7% maximum error.

**Figure 3.**
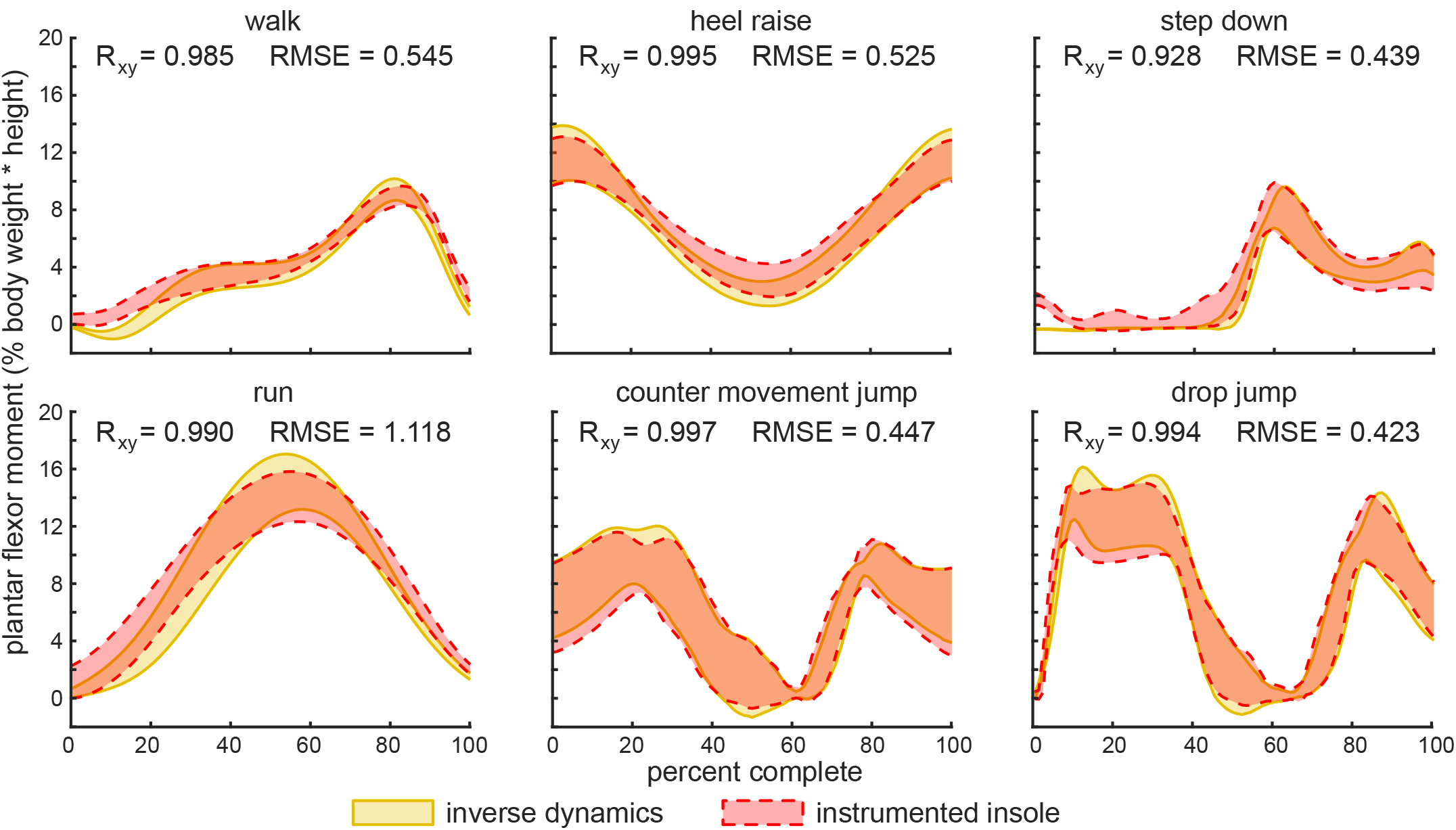
(double column): Plantar flexor loading measured by the instrumented insole (dashed red curves) was compared to inverse dynamics (solid gold curves). These values were reported as a percentage of the plantar flexor moment that we normalized by subject height and weight. Plantar flexor loading is visualized as a 95% confidence interval of the tested subjects. The waveforms were compared using calculate root mean square errors (RMSE) across each movement and shown to be strongly correlated. These RMSE values are also reported as a percentage of subject height and weight. Cross correlation coefficients also show very strong agreement between the two measures.

## Discussion

The purpose of this study was to validate a novel method for measuring plantar flexor loading outside of a traditional gait laboratory using a commercially available instrumented insole. Our findings demonstrate that simply calculating the weighted average of discretized plantar loads by sensor distances compare favorably with the motion capture (peak loading errors < 7%), the gold standard for biomechanical assessment during functional activities. While this type of instrumented insole has previously quantified loading magnitude and frequency [9–11], our study is the first to our knowledge to extend these plantar measurements to obtain motion capture quality measurements of joint level function.

Our measurements of plantar flexor loading compare favorably with reported tensiometer, inverse dynamics, and force buckle measurements during walking, running, and jumping [12–14]. This builds confidence that our method does not sacrifice measurement fidelity while being relatively low-cost and simple to implement. However, there were several limitations to consider. This approach appears better suited for quantifying plantar flexor loading than dorsiflexion loading as we did not detect dorsiflexion loading during the early weight acceptance of the gait cycle (**Figure 3**). Additionally, due to the constraints of this instrumented insole, our approach only approximates sagittal joint loading. However, a similar discretized insole is available with medial and lateral sensing zones. Lastly, we controlled the footwear worn by all research subjects (Air Pegasus, Nike, Beaverton, OR), but our preliminary data integrating this insole into an immobilizing boot suggest that different footwear does not affect measurement fidelity.

Our findings demonstrate the utility and fidelity of a new method for measuring plantar flexor loading using a commercially available instrumented insole. Quantifying plantar flexor loading is critical for understanding athletic performance [1], mobility constraints in the elderly [2], and treating Achilles tendon injuries [15]. By leveraging this new methodology, it is now feasible to prospectively track and eventually prescribe plantar flexor loading outside of the clinic to improve patient outcomes.

## Acknowledgements

the Authors have no acknowledgements

